# Polynomial Mendelian Randomization reveals widespread non-linear causal effects in the UK Biobank

**DOI:** 10.1101/2021.12.08.471751

**Authors:** Jonathan Sulc, Jennifer Sjaarda, Zoltán Kutalik

## Abstract

Causal inference is a critical step in improving our understanding of biological processes and Mendelian randomisation (MR) has emerged as one of the foremost methods to efficiently interrogate diverse hypotheses using large-scale, observational data from biobanks. Although many extensions have been developed to address the three core assumptions of MR-based causal inference (relevance, exclusion restriction, and exchangeability), most approaches implicitly assume that any putative causal effect is linear. Here we propose PolyMR, an MR-based method which provides a polynomial approximation of an (arbitrary) causal function between an exposure and an outcome. We show that this method provides accurate inference of the shape and magnitude of causal functions with greater accuracy than existing methods. We applied this method to data from the UK Biobank, testing for effects between anthropometric traits and continuous health-related phenotypes and found most of these (84%) to have causal effects which deviate significantly from linear. These deviations ranged from slight attenuation at the extremes of the exposure distribution, to large changes in the magnitude of the effect across the range of the exposure (e.g. a 1 kg/m^2^ change in BMI having stronger effects on glucose levels if the initial BMI was higher), to non-monotonic causal relationships (e.g. the effects of BMI on cholesterol forming an inverted U shape). Finally, we show that the linearity assumption of the causal effect may lead to the misinterpretation of health risks at the individual level or heterogeneous effect estimates when using cohorts with differing average exposure levels.

## 1 Introduction

Identifying factors that cause disease or influence disease progression is integral to both furthering our understanding of disease pathophysiology and helping inform the development of novel treatments and interventions. Randomized controlled trials (RCTs) are the gold standard for demonstrating such causality between an exposure and outcome, however they are expensive, time-consuming, and often unethical or infeasible. Mendelian randomization (MR) has proven to be an extremely reliable, cost-effective, and feasible alternative to RCTs to assess causality using large-scale observational genetic data. MR takes advantage of the fact that genetic variants are both determined at birth and inherited randomly and independently of other risk factors of a disease, using genetic variants as instrumental variables (IVs) to infer causality between an exposure and an outcome. This random allocation of genetic variants minimizes the possibility of reverse causality and confounding. MR has not only identified thousands of novel, causal relationships between risk factors and diseases, but also provided strong evidence for a non-causal effect of many other exposure-outcome relationships [1].

MR relies on three core assumptions: (1) relevance (i.e. IVs must be associated with the exposure), (2) exchangeability (i.e. IVs must not be associated with any confounder in the exposure-outcome relationship), and (3) exclusion restriction (i.e. IVs must not affect the outcome except through the exposure). While many extensions of MR have been developed to address potential violations of the three core assumptions, nearly all MR approaches implicitly assume that any putative causal effect is linear in nature. However, the association between an exposure and outcome is in fact often nonlinear. For instance, the observed relationship between BMI and all-cause mortality has been repeatedly shown to be U-shaped in nature, where increased mortality risk exists on either side of the 20-24.9 kg/m^2^ interval[2]. Additionally, many asymptotic relationships have been observed in the context of public health, whereby an intervention or risk factors appear to be beneficial or increase risk initially and subsequently the effect plateaus after a certain threshold [3, 4]. Along these lines, weight loss may reduce LDL levels for only lean individuals due to an inverted U-shape relationship [5]. Currently, it is unclear whether these observations are merely an artifact of confounding and/or reverse causation, or whether there is truly a causal relationship, non-linear in nature, underlying these observations.

While identifying causal relationships is of high research importance, of equal importance is a comprehensive understanding of the underlying mechanisms and dynamics. A biased and nave characterization of these causal relationships can lead to a misinformed understanding of disease mechanisms and ultimately misguided treatments and public health recommendations and interventions. To date, few approaches have been developed to investigate nonlinear causal relationships within an MR framework. Most approaches are semiparametric in nature and involve stratifying individuals based on the exposure distribution[6] or do not consider potential non-linear confounder effects[7]. The extra flexibility regarding the shape of the causal function offered by these semi-parametric methods comes at a cost of increased variance of the resulting estimator. To address these gaps, we developed PolyMR, an MR-based approach to assess non-linear causal relationships in a fully parametric fashion.

## 2 Methods

Let *X* and *Y* denote two random variables representing complex traits. We intend to use Mendelian Randomization (MR) to estimate a non-linear causal effect of *X* on *Y*. The genotype data of the SNPs to be used as instrumental variables (IVs) is denoted by *G*. To simplify notation we assume that *E*(*X*) = *E*(*Y*) = *E*(*G*) = 0 and *V ar*(*X*) = *V ar*(*Y*) = *V ar*(*G*) = 1. The effect sizes of the instruments on *X* is denoted by ***β***. Let us assume the following models

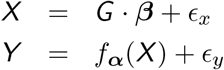

where the parametric function *f*_***α***_(·) determines the shape of the causal relationship between *X* and *Y* and *E*_*x*_ and *E*_*y*_ are zero-mean errors. For simplicity, we assume that *f*_***α***_(·) is a polynomial – even if it is not it can be approximated by one with arbitrary precision over the range of the majority of values *X* can take. For example, if we intend to test a quadratic causal relationship, *f*_***α***_(*x*) = *α*_0_ + *α*_1_ · *x* + *α*_2_ · *x*^2^. Thus, the model can be rewritten as

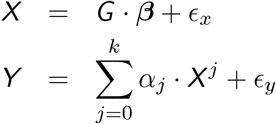

The above equation can be expanded to

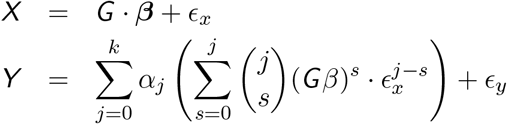

We rely on the INSIDE assumption [8], which ensures that *cov*(*Gβ, ϵ*_*y*_) = 0. The error terms *ϵ*_*x*_ and *ϵ*_*y*_ can nevertheless be correlated because of a potential causal effect of *Y* on *X* (reverse causation) and/or due to confounders. Let us split *ϵ*_*y*_ into *ϵ*_*x*_-dependent and *ϵ*_*x*_-independent parts.

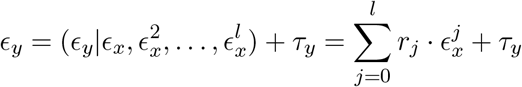

Since *cov*(*G****β***, *ϵ*_*x*_) = *cov*(*G****β***, *ϵ*_*y*_) = 0, the residual noise *τ*_*y*_ is independent of both *G****β*** and *ϵ*_*x*_. As a consequence *cov*(*X, τ*_*y*_) = *cov*(*X* − *G****β***, *τ*_*y*_) = 0. This allows us to rewrite the main model equations

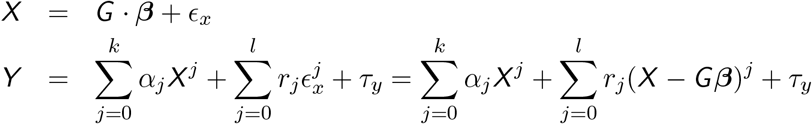

The advantage of this equation system is that the error terms (*E*_*x*_ and *τ*_*y*_) are uncorrelated and independent of the respective explanatory variables.

Let the realizations of the random variables *X, Y, G* be denoted by ***x, y***, *G*, observed in a sample of size *n*. The parameters {***β, α, r***} can be estimated by computing two ordinary least squares estimates, first estimating ***β*** using the first equation and substituting this into the second equation:

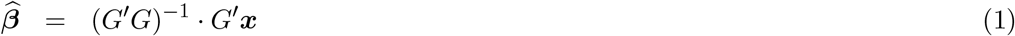

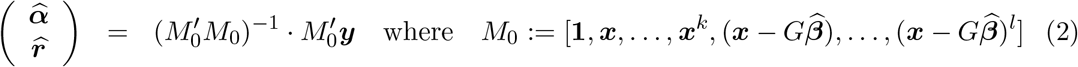

The special case of this approach when *k* = *l* = 1 is equivalent to the standard control function approach. For higher orders, the terms *k* and *l* are not required to be equal, as *k* represents the powers of the *f*_*α*_(·) function describing the causal relationship between *X* and *Y*, whereas *l* concerns the order of the confounding/reverse causation.

### 2.1 Implementation

We implemented this method in R. For the polynomial approximation of *f*_*α*_(·) function of the causal relationship, we used *k* = 10. We then iteratively eliminated coefficients which were not significant at a Bonferroni-corrected level (i.e. 0.05*/a*, where a is the number of coefficients remaining) by setting them to zero. We set *l* to be equal to the polynomial order of the function, retaining all terms up to *l* such that any association with higher orders of the exposure driven by confounding are properly accounted for. Once all remaining coefficients were significant, the non-linearity p-value was obtained with a likelihood ratio test (LRT), comparing the full model with that including only the linear effect but retaining all residual correction terms (*r*_*j*_). The causally explained variance was determined as the difference in explained variance (*r*^2^) between the full model and that excluding all *α*_*j*_ · ***x***_*j*_ terms (i.e. accounting only for potential confounding).

The polynomial function was the result of a multivariable regression, which also provided us with the variance-covariance matrix of the coefficient estimates. From these, we can generate causal polynomial functions whose coefficients are drawn from the established multivariable distribution to obtain the 95% confidence hull.

In order to avoid invalid IVs acting through reverse causation, we filtered out IVs where the standardized effect estimate was larger (in absolute value) in the outcome than in the exposure. For comparison, LACE [6] was also implemented and polynomial approximation was obtained in an analogous fashion. The piece-wise linear LACE approach was not tested here but is considered in the Discussion. We examined the limitations of the standard (1st order) control function approach by running PolyMR with *l* = 1, hereafter referred to as PolyMR-L1, in specific settings. We chose not to compare with results from CATE [7] as this method enables the estimation of differences in causal effects based on exposure (i.e. slopes) but does not provide an overall function and makes comparison difficult. Furthermore, the implementation available at the time of this writing was comparatively slow and did not scale well when tested on biobank-size data: the estimated run-time for a single simulation using our parameters, repeated at ∼100 exposure levels to model function shape, was several days for fewer than 10 IVs.

### 2.2 Simulations

We simulated data according to the following model

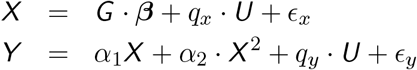

where *U* is a confounder drawn from a standard normal distribution. Columns of *G* were drawn from a binomial distribution with minor allele frequencies following a beta distribution with shape parameters equal to 1 and 3, and then normalized each column to have zero mean and unit variance. The genetic effects *β*_*i*_ were drawn from normal distributions based on the minor allele frequencies, specifically *β*_*i*_ ∼ 𝒩 (0, (*p*_*i*_ * (1 − *p*_*i*_))^−0.25^), where *p*_*i*_ is the minor allele frequency of SNP i, and scaled such that the total explained variance matches the predefined heritability, i.e. 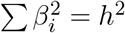. These effect sizes are realistic and are according to the baseline LDAK heritability model (without functional categories) with selection strength of −0.25 [9, 10]. For the basic settings, we included moderate confounding (*q*_*x*_ = 0.2 and *q*_*y*_ = 0.5) and a quadratic causal function (*f*_***α***_(*X*) = 0.1*X* + 0.05*X* ^2^). The heritability *h*^2^ was set to 0.5, explained by *m* = 100 causal SNPs, and sample size was set to 100,000 individuals. Causal SNPs were filtered for genome-wide significance of their marginal effects in the simulated data prior to their use as IVs. Variations on these settings were tested, as shown in table 1. Each combination of parameters was used to generate 1000 sets of data, to which we applied both PolyMR and LACE. We compared these with the performance of PolyMR-L1 in the base settings, in the presence of weak quadratic confounding (*q*_*x*_ × *q*_*y*2_ = 0.04), as well as in the absence of quadratic causal effect but with quadratic confounding (*q*_*x*_ × *q*_*y*2_ = 0.1) creating a similar observed association between traits as in the base settings.

**Table 1:**
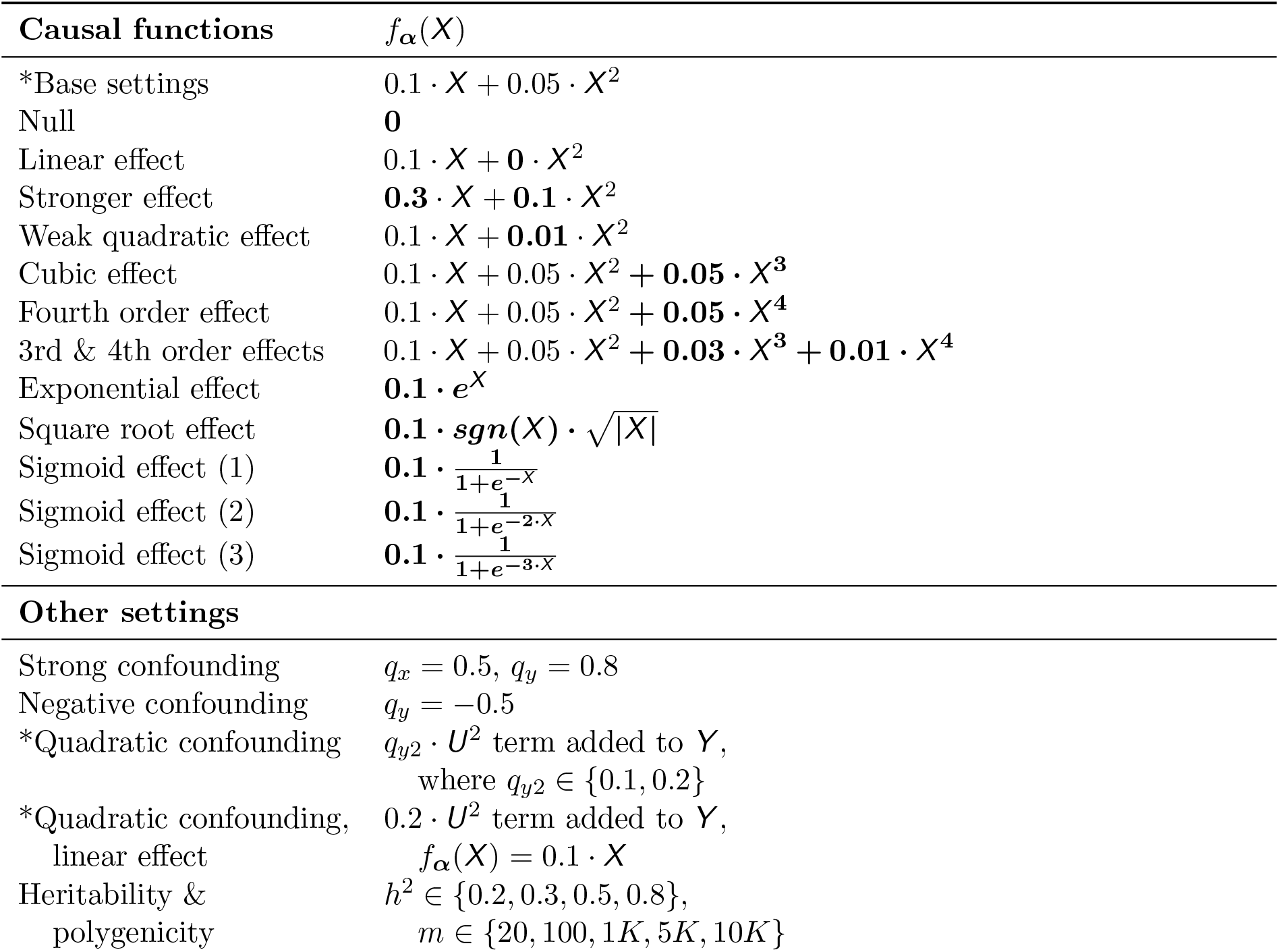
Causal functions and setting parameter combinations. simulated for PolyMR. Modifications to the base setting’s causal function are indicated in **bold**. Asterisks (*) denote settings where PolyMR-L1 was also applied for comparison.

The theoretical 95% confidence hulls were compared to the empirical distribution of estimated models. At each percentile of the exposure distribution, the size of the predicted 95% confidence intervals (CIs) was compared to the empirical one.

### 2.3 Application to UK Biobank data

The UK Biobank is a prospective cohort of over 500,000 participants recruited in 2006–2010 and aged 40–69 [11]. We tested for non-linear causal effects of anthropometric traits (body mass index [BMI], weight, body fat percentage [BFP], and waist-to-hip ratio [WHR]) on continuous health outcomes (pulse rate [PR], systolic blood pressure [SBP], diastolic blood pressure [DBP], and glucose, and LDL, HDL, and total cholesterol [TC] levels in blood), as well as the reverse. We also tested for effects of both BMI and age completed full time education on life expectancy. Since most participants in the UK Biobank are still alive, we scaled the mother’s and father’s age of death (separately) and used the mean of these standardized phenotypes as a proxy for the individual’s life expectancy.

We selected 377,607 unrelated white British participants and all phenotypes were corrected for age, age^2^, sex, age × sex, age^2^ × sex, as well as the top 10 genetic principal components. With the exception of WHR, IVs were selected using the TwoSampleMR R package [version 0.5.5, 12] with default settings (*p* < 5 * 10^−8^, *r*^2^ < 10^−3^, *d* > 10^4^ kb) from the GWAS in the ieugwasr R package [version 0.1.5, 13] with the largest number of instruments overlapping our dataset. For WHR, we used a previously-performed GWAS on the aforementioned sample from the UK Biobank, adjusting for covariates as above [14].

For the purpose of comparison, we also used inverse-variance weighted MR and MR Egger on each of these exposure-outcome pairs. These were performed with the same IVs and (in-sample) association statistics using the TwoSampleMR R package [version 0.5.5, 12]. We also compared the results of standard PolyMR with those PolyMR-L1 to determine whether accounting for higher order confounding is necessary in real data applications.

## 3 Results

### 3.1 Simulations

We simulated a variety of settings, including many combinations of heritability and polygenicity in the exposure, sample size, and shape of the causal function *f*_***α***_(·) and confounding. Where the true underlying function was polynomial, our approach correctly captured its shape (Fig. 1) although a slight bias from confounding was introduced in certain settings with high polygenicity (*>* 1000 causal SNPs) or strong confounding (where linear confounding explained ∼40% of the exposure-outcome association) (Fig. 2). The distribution of this bias was affected by the shape of the confounding, i.e. in situations with quadratic confounding, the bias was quadratic with respect to exposure. In all simulation settings, this bias was orders of magnitude smaller than both the causal effect and the confounding (e.g. Fig. 1A). Although this bias was minimal with the standard PolyMR settings (*l* = *k*), quadratic confounding produced significant bias when higher orders of the control function term 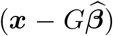 were ignored (i.e. *l* = 1 in PolyMR-L1, Supplementary Figure 1), which is the standard approach for control function use.

**Figure 1:**
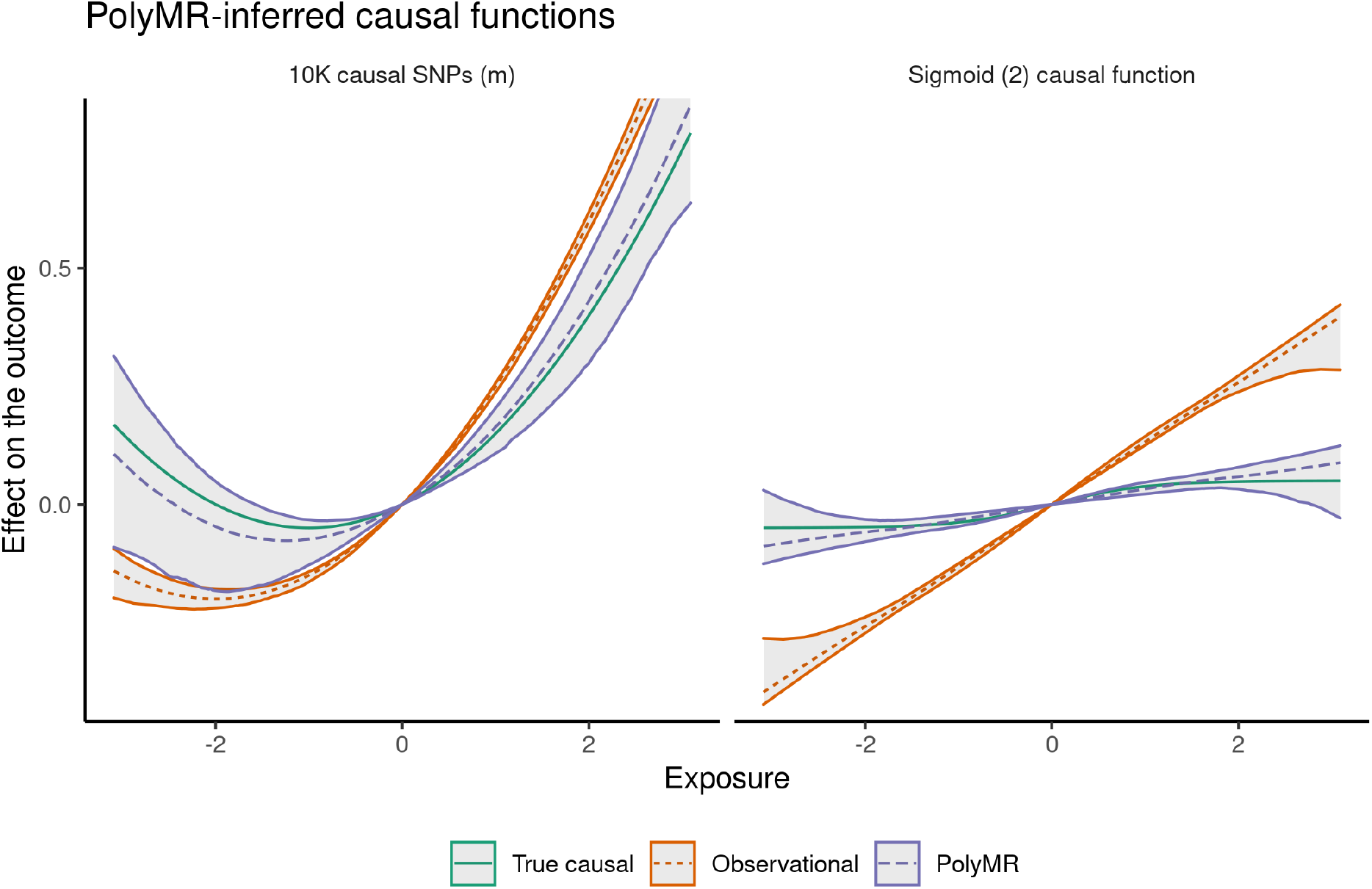
PolyMR is able to recover the shape of the causal function. The true causal function is shown in green (solid line). The observed association model is shown in orange (short-dashed) while that obtained using PolyMR is shown in purple (long-dashed). The hulls around the model curves show the 95% coverage hull across 1000 simulations. The settings shown here represent high polygenicity (10’000 causal SNPs accounting for a heritability of 0.3 - left panel); and a sigmoid causal effect 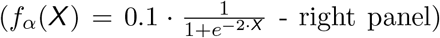 scenarios. The Y-axis shows the expected association with/effect of the exposure on the outcome, relative to the outcome level at the mean population exposure.

**Figure 2:**
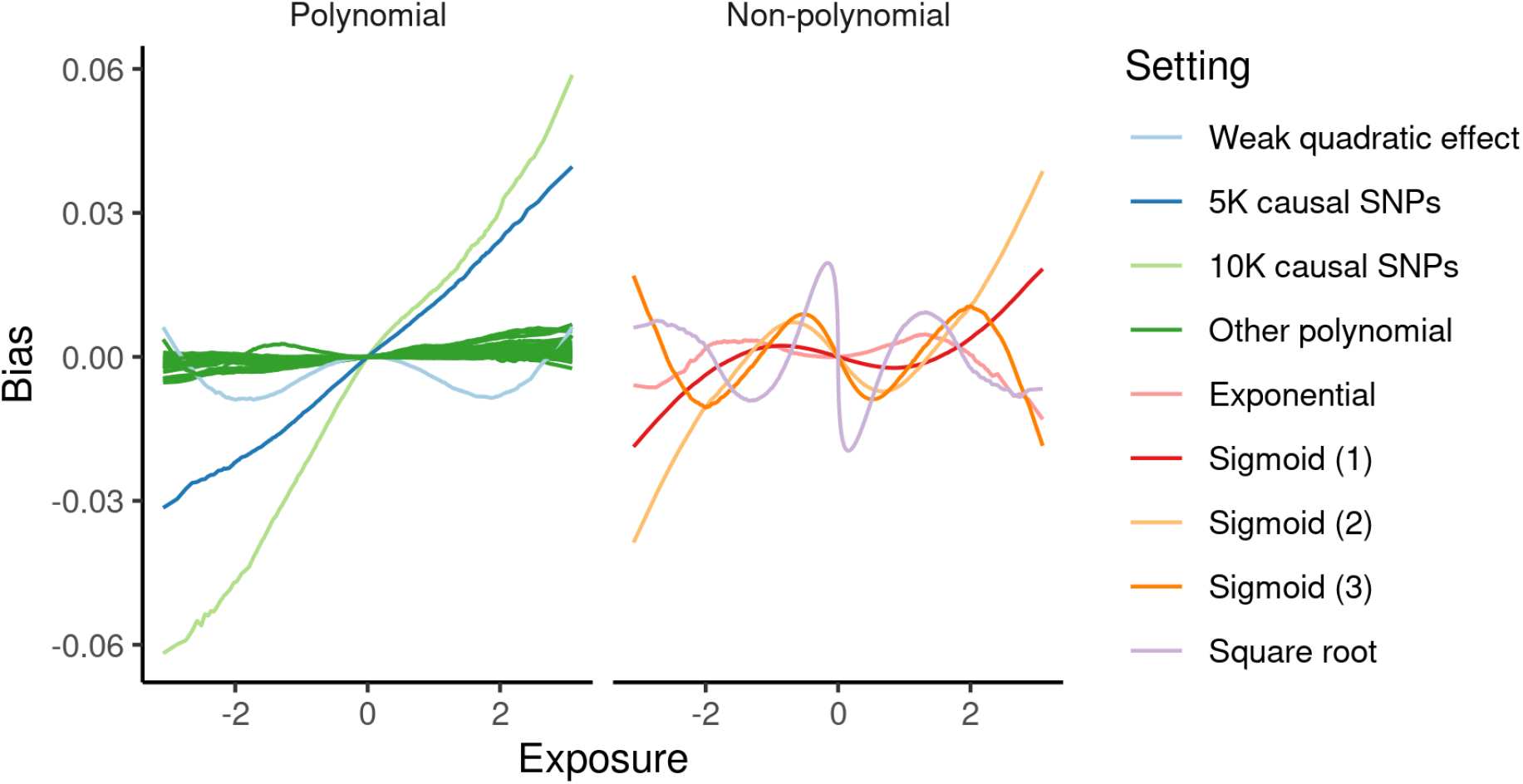
Bias as a function of exposure across settings. In settings with a polynomial causal function *f*_***α***_(·), slight bias from non-genetic confounding was induced under certain combinations of high polygenicity, high heritability, or strong confounding. The bias found in non-polynomial settings was expected due to the polynomial approximation approach.

In the case of non-polynomial functions, PolyMR nevertheless provided reasonable estimates of the true shape of the causal function (Fig. 1B). The bias introduced in these cases (Fig. 1B, 2B) is consistent with expectations of polynomial approximation with limited power and dependent on the shape of the non-polynomial function.

LACE also produced some bias in estimating the causal function. The magnitude of the bias introduced by either method was dependent on the settings used. Still, in all settings tested, PolyMR provided lower bias (Supplementary Figure 2) and root mean square errors (RMSEs) than LACE (Fig. 3), partly driven by greater statistical power and smaller SEs.

**Figure 3:**
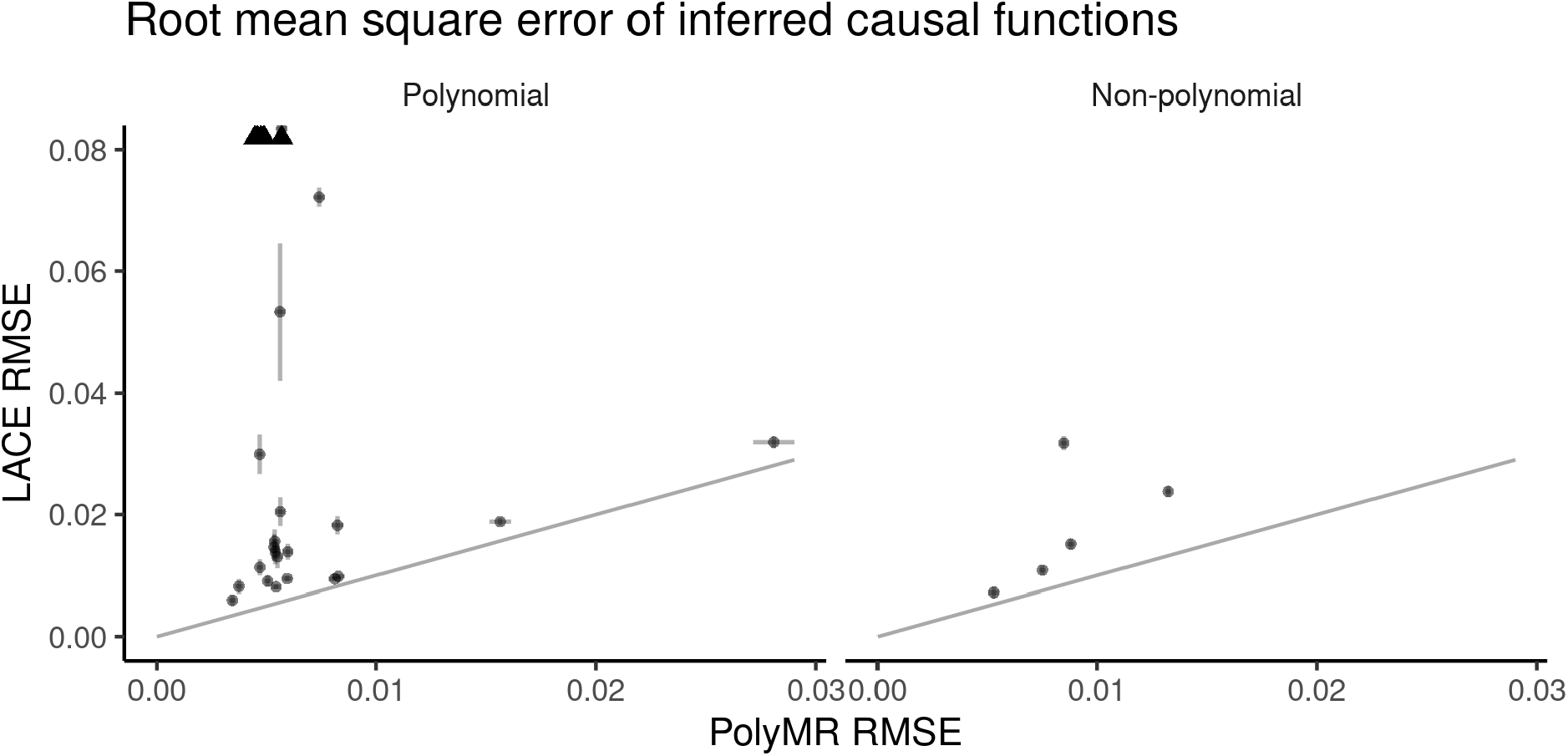
PolyMR provided greater accuracy in the estimation of causal functions. The root mean square errors (RMSEs) are shown for both PolyMR and LACE. Each point is the mean RMSE for a given setting with the error bars showing the 95% confidence interval of the mean. Settings were split into polynomial or non-polynomial causal functions. Arrows in the polynomial plot indicate RMSEs which exceed the bounds of the plot.

To ensure that the variance estimated from the variance-covariance matrix of the model was correctly calibrated, we assessed the coverage of the 95% confidence interval. We did so by comparing the predicted 95% confidence intervals (CIs) of the curves with those derived empirically from repeated simulations. We found that in the case of most polynomial functions, the CIs were properly calibrated with the theoretical and empirical CIs being almost equal across most of the exposure distribution. Note that under some simulation settings (e.g. weak quadratic effects, *α*_2_ = 0.01) allowing the polynomial degree to vary led to increased empirical variance (see Supplementary Fig. 3A). However, if we consider only those simulations where the second order was correctly inferred (920 simulation results out of 1000) the empirical CIs were close to those predicted by the method (Supplementary Fig. 3B).

### 3.2 UK Biobank

Given its favorable performance throughout all simulation settings, we applied the PolyMR method to data from the UK Biobank. We set out to estimate the causal effects of four anthropometric traits (BMI, weight, BFP, and WHR) on each of seven continuous traits commonly used as health biomarkers (SBP, DBP, PR, and the levels of glucose, HDL, LDL, and total cholesterol in the blood). We also tested for reverse causal effects for these trait pairs, as well as any effects of BMI or education on life expectancy.

The effects of the anthropometric traits were qualitatively similar to one another and significant against all tested outcomes with significant non-linearity in most cases (Supplementary Figures 4–31). Those of BFP and WHR tended to be more similar to one another, monotonically increasing DBP, SBP, and PR with linear to slightly non-linear effects. BMI also increased these traits overall, though the effects of BMI on DBP and SBP plateaued at around 2 SD above the population mean (∼36.9 kg/m^2^) and the causal function for BMI on PR showed a positive slope for values between approximately 1 SD below to 2 SDs above the population mean (∼22.7–36.9 kg/m^2^) with negative slopes beyond these. The effects of weight were weaker but qualitatively similar to those of BMI. Glucose was increased by all of these, though the effects of a change in exposure were negligible below ∼-1 SD for all traits and intensified at higher values. For example, the estimated slope of the standardized effect of BMI was 0.17 at the population mean but increased to 0.31 at +2 SD. The strongest non-linearity in the effects of anthropometric traits was found for total cholesterol, mainly driven by the LDL fraction (Fig. 4A), where the causal function took a strong inverted U-shape. In contrast to this, their effects on HDL were all monotonic decreasing.

**Figure 4:**
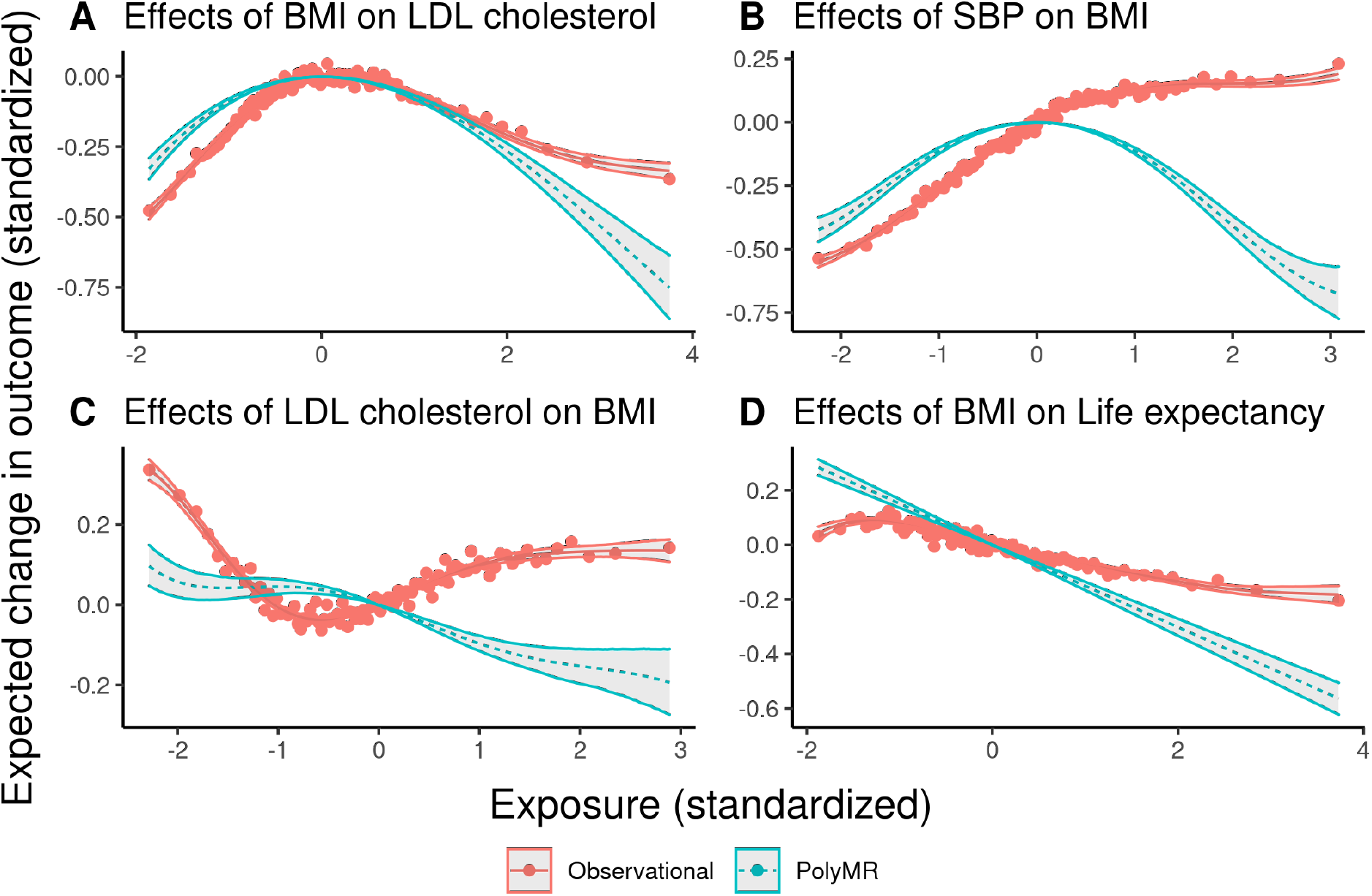
Most tested causal effects have strong non-linear components in the UK Biobank. The red points show the mean outcome plotted against the median exposure for each of 100 bins, split by covariate-adjusted exposure level. The red curve (solid) is the multivariable regression model whereas the teal one (dashed) corresponds to the estimated causal function obtained using PolyMR. The hulls around both curves correspond to the 95% confidence interval.

The consequences of PR were limited to weak linear effects on BFP and WHR. TC linearly decreased BMI, weight, and BFP, with no detectable effect on WHR. SBP and DBP both had inverted U-shaped causal functions for their effects on all anthropometric traits (*p* < 1.6*10^−47^), with the effects on WHR being slightly weaker. Glucose levels had nearly no effect on most traits across most of the distribution, but drove strong reductions at higher values, with the exception of WHR, which was in fact slightly increased by glucose levels up to ∼ 3 SD before being decreased at higher levels. HDL had a slight U-shaped effect on these traits, with a stronger increase for high values of the exposure on BFP and no increase in WHR.

Although the observational association of SBP/DBP and the anthropometric traits was mostly monotonic increasing, the estimated causal function on these had an inverted U-shape (4B), with slightly weaker effects on WHR. The causal effects of PR on anthropometric traits show a slight positive slope close to the population median, but the directionality switches at either extreme of the distribution. LDL cholesterol decreased the outcomes near monotonically, though the effect close to the population median was weak to null (Fig. 4C). The impact of glucose was slightly different across the anthropometric traits. Both BMI and weight were overall negatively affected by glucose levels (with weaker effects around zero). BFP was also decreased, although the effect was much weaker. WHR, however, was slightly increased by glucose levels up to ∼3 SD before being decreased at higher levels. Note that the effect close to the population mean is likely driven by a decrease in hip circumference rather than an increase in the waist’s, similar to what we’ve shown previously for the effects of diabetes risk and triglyceride levels on WHR-related metrics [14].

The effects of BMI and education on life expectancy are directionally as expected but we found no evidence of non-linearity. The BMI-life expectancy (causal) relationship was decreasing, though the intensity of the effect was greater than the observed association (Fig. 4D). As expected, higher education increased life expectancy but we found no evidence of non-linearity in the effect (Supplementary Figure 32).

The exclusion of higher order control function terms in PolyMR-L1 produced somewhat different inferred causal functions, with generally stronger non-linear components, resulting in inferred causal functions which were closer to the observed associations (Supplementary Figures 33–39).

## 4 Discussion

In this report, we present PolyMR, a Mendelian randomization-based approach for the inference of non-linear causal effects. Through a variety of simulations, we showed that it is robust to many forms of confounding and is well-powered to detect even weak quadratic effects in biobank-size cohorts. Finally, by applying our method to the UK biobank, we showed that causal effects across many anthropometric traits indeed include strong non-linear components.

Despite statistically significant non-linearity for the causal effects of many exposures on outcomes, some of these were monotonic or even near linear around the population median. In these cases, the causal effect estimates from traditional, linear, MR methods will likely still be useful, though non-linearity in the tails of the distribution may introduce varying amounts of bias for certain exposure levels. Even where the overall causal effect is monotonic and could therefore be described as “positive” or “negative,” knowing the shape of the curve can provide insight into which strata of the population may benefit most from public health interventions. For example, weight loss for individuals with average BMI is far more beneficial in in terms of lowering systolic blood pressure than it is for obese individuals. The bias introduced by the assumption of linearity increases further with non-monotonic effects, such as that of SBP on BMI, and the effect estimates vary greatly based on the method used, but without proper consideration of the non-linear components will not be particularly meaningful. Although in certain contexts, a linear approximation of the monotonic effects can be useful, they will introduce bias. Specifically, different populations with different mean values of exposure will yield different linear estimates even in cases where there is true causality and all IVs are valid.

There are a number of possible explanations for non-linear causal relationships we identified, particularly in the case of the inverted U-shaped curves, where both the exposure and outcome are presumed negative markers of health (for example BMI and cholesterol). The shape of these effects could arise from several mechanisms such as negative feedback loops. The most obvious, but intangible, candidates are biological feedback mechanisms. A more concrete possibility is a lifestyle change in response to elevated risk, by either doctor recommendation, medication, or personal or social pressures. Either off- or on-target effects of these changes could play a role on the inverted U-shaped curves we identified. These latter effects, however, are expected to be weak and explain only a small part of these phenomena. Another explanation may be interaction effects between an exposure-associated environmental variable and the exposure itself influencing the outcome.

We identified several benefits of our method compared to existing tools, such as LACE and CATE. PolyMR not only demonstrated greater accuracy than LACE, but it does not require arbitrary choices regarding bin numbers and spacing, like LACE does. The other competing method, CATE, assumes that any sources of confounding are linear and will introduce bias/false positives in the presence of sources of non-linear confounding, whereas polyMR allows for non-linear sources of confounding. Of note, while CATE can estimate the change in outcome based on the difference in exposure, it does not explicitly estimate the shape of the causal function.

This work has certain limitations that should be taken into account. First, our method requires individual-level data. While using classical summary statistics, non-linear causal effects are undetectable, however one could envision approximating the polyMR method by using not only *G* − *X* and *G* − *Y* association summary statistics but higher order (*G*^*i*^ − *Y, G*^*i*^ − *X*^*j*^, *X*^*i*^ − *Y* and *G*^*i*^ · *X*^*j*^ − *Y* with *i, j* = 0, 1, 2, …, *k*) associations. Although such an approximation would require additional summary statistics, those are not generally available for any trait, hence would not facilitate its use in practice. Secondly, the estimates provided for the extremes of the distribution are less reliable, and removing outliers may improve the reliability of estimates across the entire distribution. Third, a small amount of bias is introduced due to the fact that we use 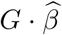 instead of *G* · *β* in our model fitting (Eq. (2)). To mitigate this, MLE could also be used to take into account the error in SNP-exposure associations. Fourth, our approach still suffers from the weaknesses of classical MR methods, such as Winner’s curse and invalidity of the instruments. Finally, although this method could be generalized to binary outcomes, it has limited utility. The shape of such non-linear relationships would largely depend on the link function used for the generalized linear models. While showing deviations from a linear relationship for continuous outcomes reveals not only a quantitatively better fitting model, but also qualitatively different. However, for binary outcomes, even if there is a non-linear term, the model class remains qualitatively similar (since any link function is already non-linear) and it only indicates that the causal relationship could be better described with a different kind of link function. The only exception from this is when the causal function is non-monotonic, since link functions are strictly monotonically increasing. Therefore, the method is best suited to continuous traits, but may also reveal interesting insights for non-monotonic causal functions for binary outcomes.

In summary, we have developed a novel MR approach for the estimation of nonlinear exposure-outcome causal effects. We have shown the utility of this approach when applied to cardiovascular and anthropometric traits in the UK Biobank, where we identified numerous relationships that show significant deviations from linearity. Indeed, non-linear effects are pervasive in biology and should be considered appropriately when developing public health policies. Future studies should investigate the impact of non-linear causal effects on the complete human phenome to determine the prevalence of non-linear causal relationships among other conditions and biological pathways. PolyMR allows for more precise understanding of causal mechanisms beyond mere identification of causal factors for disease. Better understanding of such complex, non-linear relationships will help make better-informed health interventions.

## Data availability

The UK Biobank summary statistics data used in this study came from the Neale Lab and can be downloaded from http://www.nealelab.is/uk-biobank. UK Biobank data are available through a procedure described at http://www.ukbiobank.ac.uk/using-the-resource/.

## Code availability

The source code for this work can be found on https://github.com/JonSulc/PolyMR.

## Acknowledgements

This research has been conducted using the UK Biobank resource (#16389), which has been approved by the National Research Ethics Service Committee. The computations have been carried out on the HPC server of the Lausanne University Hospital. Z.K. was funded by the Swiss National Science Foundation (31003A-143914, 310030-189147).

## Author contributions

Z.K. and Jo.S. conceived the method. Jo.S. performed the simulation studies, the analysis of real data and wrote the initial draft of the paper with contributions from Je.S. and Z.K. All authors have read and approved the manuscript.

