## Supplementary Information for "Polynomial Mendelian Randomization reveals widespread non-linear causal effects in the UK Biobank"

### Supplementary figures

November 26, 2021

### 1 Simulations

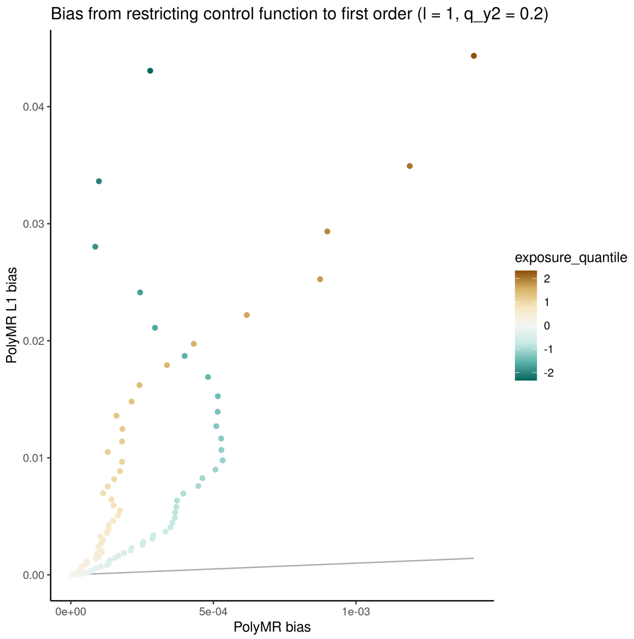

**Supplementary figure 1: Not accounting for higher order confounding introduces bias.** The (absolute) biases are shown for both PolyMR and PolyMR-L1 in the setting with quadratic confounding. Each point is the mean absolute bias across simulations for a given percentile of the exposure distribution.

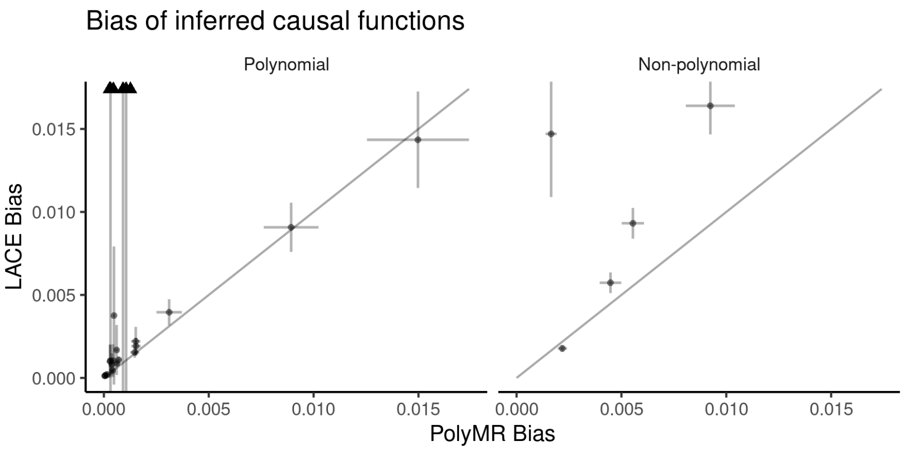

**Supplementary figure 2: PolyMR generally yields less bias than LACE.** The (absolute) biases are shown for both PolyMR and LACE. Each point is the mean absolute bias across simulations for a given setting with the error bars showing the 95% confidence interval of the mean. Settings were split into polynomial or non-polynomial causal functions. Arrows in the polynomial plot indicate mean biases exceeding the bounds of the plot.

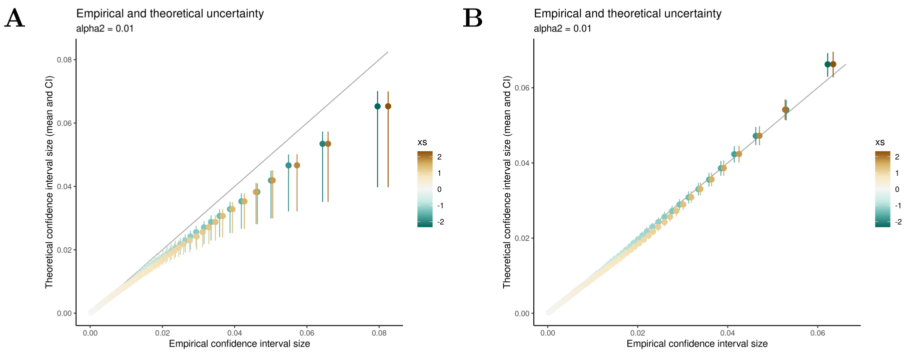

**Supplementary figure 3: The confidence intervals (CIs) of PolyMR are correctly calibrated** for the given model but may be slightly underestimated due to post-selection inference if the model is allowed to vary. The size of the theoretical 95% CI was calculated for each simulation at each percentile of the exposure, yielding a distribution of CI sizes. The empirical CIs come from the distribution of coefficient estimates across simulations. Both panels show the CIs for the weak quadratic effect setting, with either (A) all results or (B) only those results where the correct order of the causal function was determined (920 out of 1000 simulations).

### 2 Application in the UK Biobank

#### 2.1 Body fat percentage

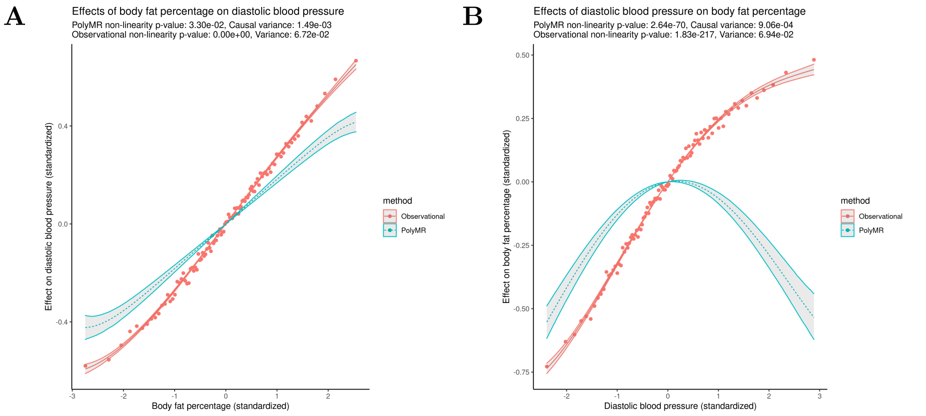

**Supplementary figure 4: The observed (red, continuous) and causal (blue, dashed) association between body fat percentage and diastolic blood pressure (A) and the reverse (B).** The observed association (red, continuous) was estimated using a similar polynomial approximation for the purpose of comparison, the points showing the mean outcome level at each percentile of the exposure. The hulls surrounding both functions show the 95% confidence range across the distribution.

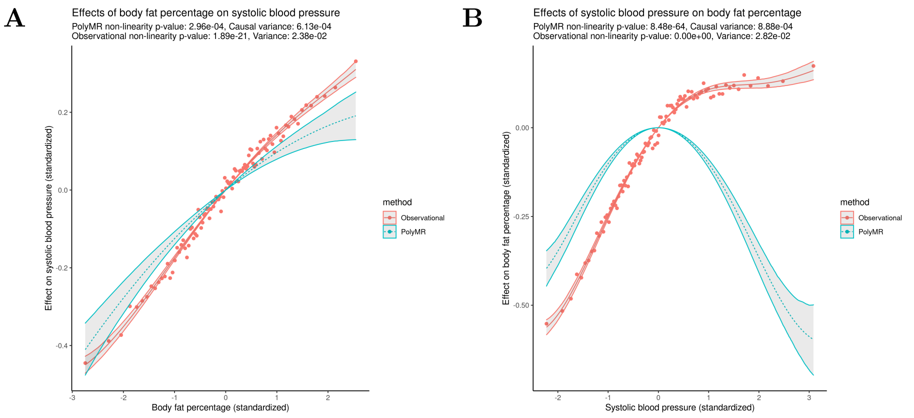

**Supplementary figure 5: The observed (red, continuous) and causal (blue, dashed) association between body fat percentage and systolic blood pressure (A) and the reverse (B).** The observed association (red, continuous) was estimated using a similar polynomial approximation for the purpose of comparison, the points showing the mean outcome level at each percentile of the exposure. The hulls surrounding both functions show the 95% confidence range across the distribution.

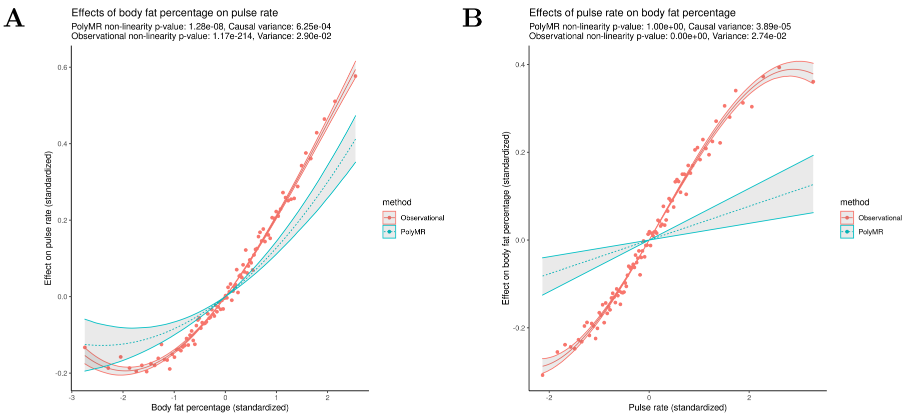

**Supplementary figure 6: The observed (red, continuous) and causal (blue, dashed) association between body fat percentage and pulse rate (A) and the reverse (B).** The observed association (red, continuous) was estimated using a similar polynomial approximation for the purpose of comparison, the points showing the mean outcome level at each percentile of the exposure. The hulls surrounding both functions show the 95% confidence range across the distribution.

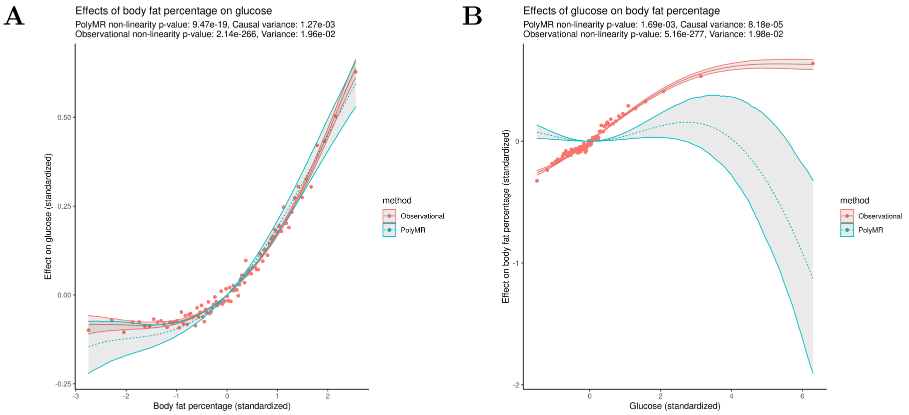

**Supplementary figure 7: The observed (red, continuous) and causal (blue, dashed) association between body fat percentage and glucose (A) and the reverse (B).** The observed association (red, continuous) was estimated using a similar polynomial approximation for the purpose of comparison, the points showing the mean outcome level at each percentile of the exposure. The hulls surrounding both functions show the 95% confidence range across the distribution.

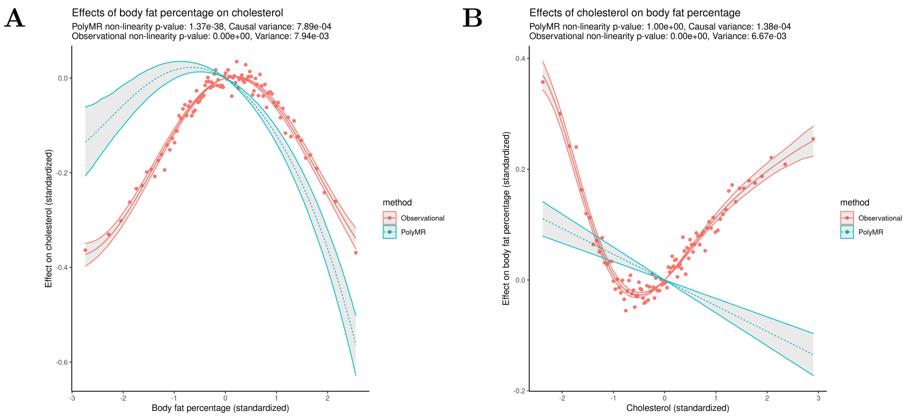

**Supplementary figure 8: The observed (red, continuous) and causal (blue, dashed) association between body fat percentage and total cholesterol (A) and the reverse (B).** The observed association (red, continuous) was estimated using a similar polynomial approximation for the purpose of comparison, the points showing the mean outcome level at each percentile of the exposure. The hulls surrounding both functions show the 95% confidence range across the distribution.

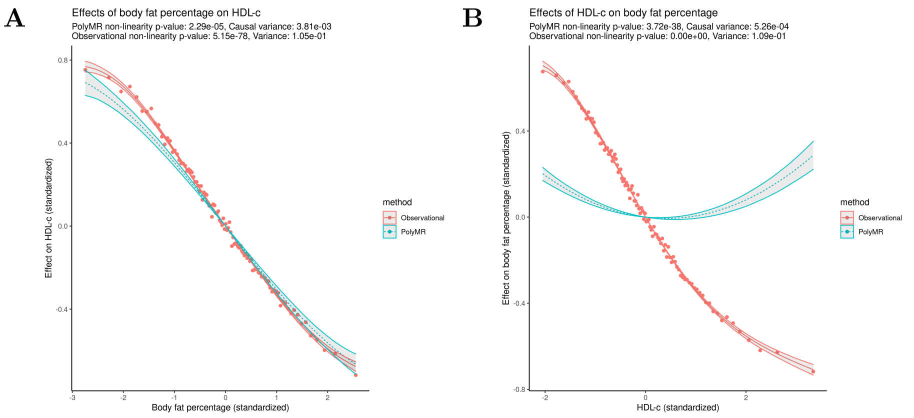

**Supplementary figure 9: The observed (red, continuous) and causal (blue, dashed) association between body fat percentage and HDL cholesterol (A) and the reverse (B).** The observed association (red, continuous) was estimated using a similar polynomial approximation for the purpose of comparison, the points showing the mean outcome level at each percentile of the exposure. The hulls surrounding both functions show the 95% confidence range across the distribution.

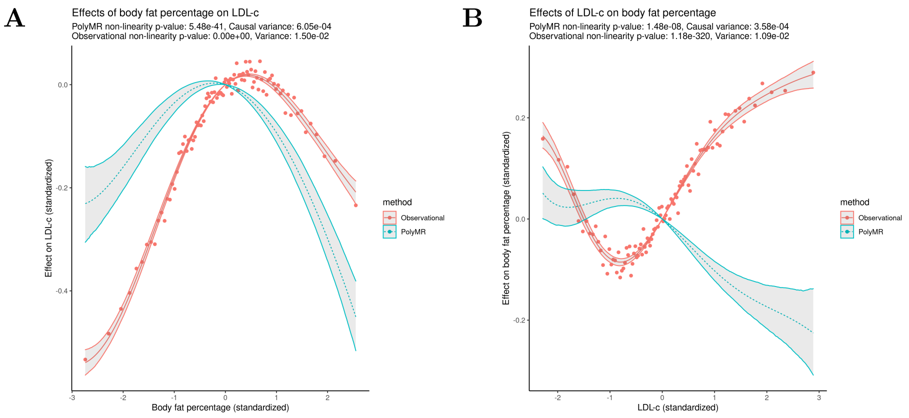

**Supplementary figure 10: The observed (red, continuous) and causal (blue, dashed) association between body fat percentage and LDL cholesterol (A) and the reverse (B).** The observed association (red, continuous) was estimated using a similar polynomial approximation for the purpose of comparison, the points showing the mean outcome level at each percentile of the exposure. The hulls surrounding both functions show the 95% confidence range across the distribution.

#### 2.2 Waist-to-hip ratio

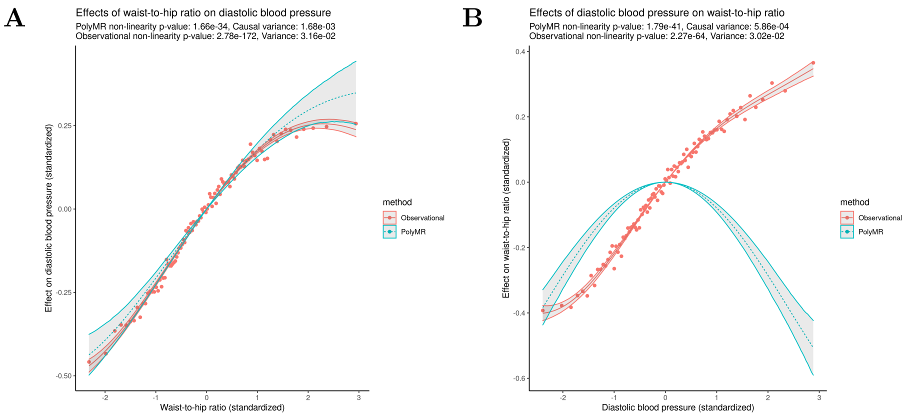

**Supplementary figure 11: The observed (red, continuous) and causal (blue, dashed) association between waist-to-hip ratio and diastolic blood pressure (A) and the reverse (B).** The observed association (red, continuous) was estimated using a similar polynomial approximation for the purpose of comparison, the points showing the mean outcome level at each percentile of the exposure. The hulls surrounding both functions show the 95% confidence range across the distribution.

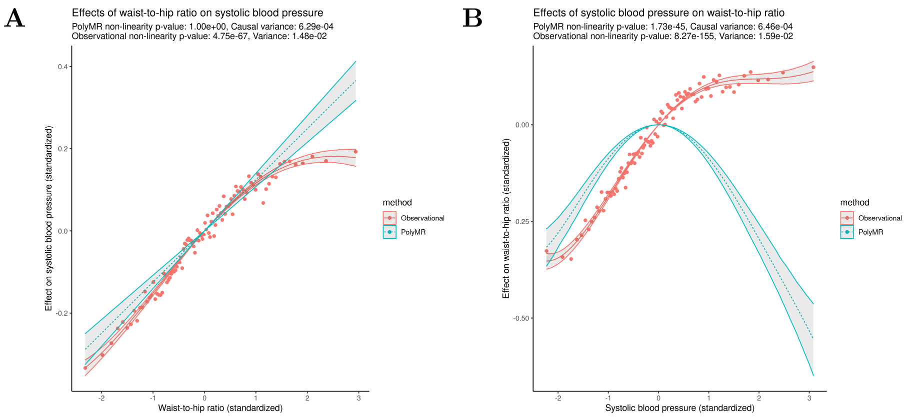

**Supplementary figure 12: The observed (red, continuous) and causal (blue, dashed) association between waist-to-hip ratio and systolic blood pressure (A) and the reverse (B).** The observed association (red, continuous) was estimated using a similar polynomial approximation for the purpose of comparison, the points showing the mean outcome level at each percentile of the exposure. The hulls surrounding both functions show the 95% confidence range across the distribution.

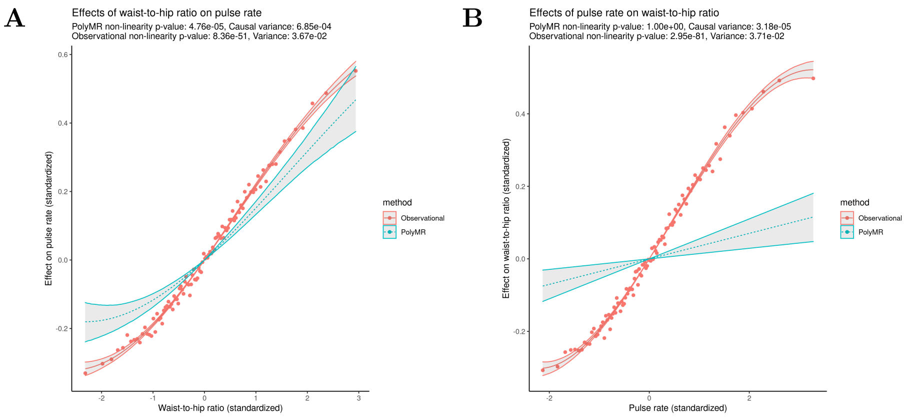

**Supplementary figure 13: The observed (red, continuous) and causal (blue, dashed) association between waist-to-hip ratio and pulse rate (A) and the reverse (B).** The observed association (red, continuous) was estimated using a similar polynomial approximation for the purpose of comparison, the points showing the mean outcome level at each percentile of the exposure. The hulls surrounding both functions show the 95% confidence range across the distribution.

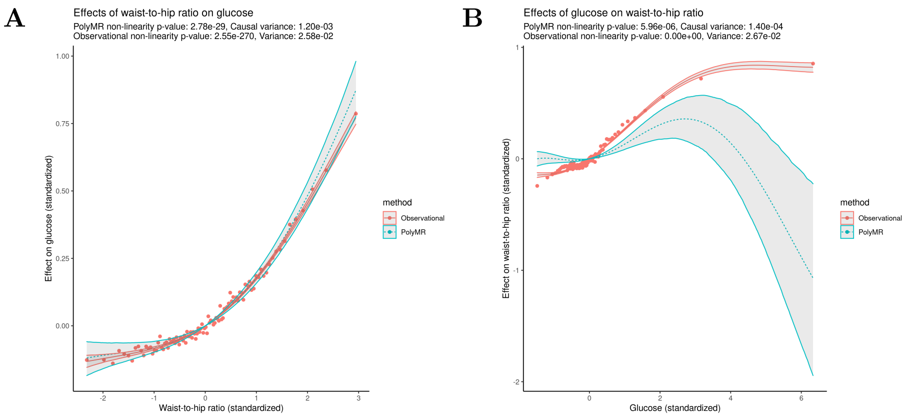

**Supplementary figure 14: The observed (red, continuous) and causal (blue, dashed) association between waist-to-hip ratio and glucose (A) and the reverse (B).** The observed association (red, continuous) was estimated using a similar polynomial approximation for the purpose of comparison, the points showing the mean outcome level at each percentile of the exposure. The hulls surrounding both functions show the 95% confidence range across the distribution.

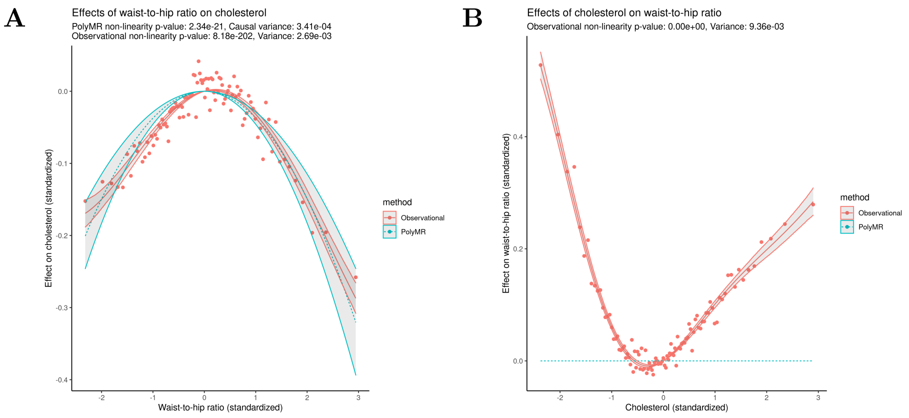

**Supplementary figure 15: The observed (red, continuous) and causal (blue, dashed) association between waist-to-hip ratio and total cholesterol (A) and the reverse (B).** The observed association (red, continuous) was estimated using a similar polynomial approximation for the purpose of comparison, the points showing the mean outcome level at each percentile of the exposure. The hulls surrounding both functions show the 95% confidence range across the distribution.

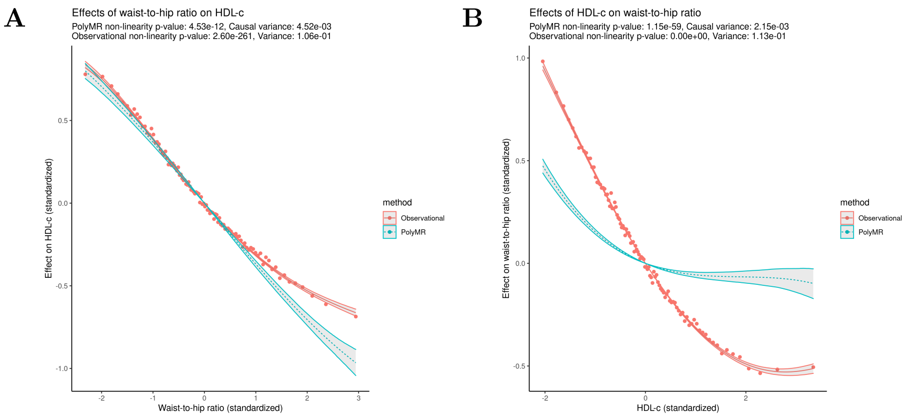

**Supplementary figure 16: The observed (red, continuous) and causal (blue, dashed) association between waist-to-hip ratio and HDL cholesterol (A) and the reverse (B).** The observed association (red, continuous) was estimated using a similar polynomial approximation for the purpose of comparison, the points showing the mean outcome level at each percentile of the exposure. The hulls surrounding both functions show the 95% confidence range across the distribution.

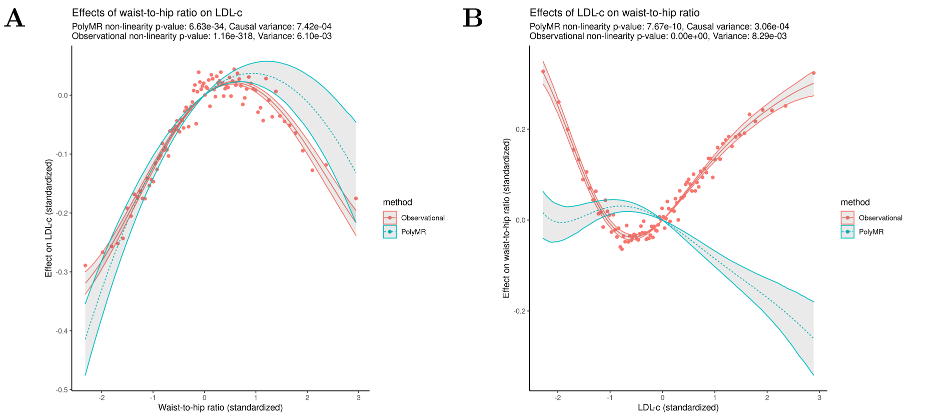

**Supplementary figure 17: The observed (red, continuous) and causal (blue, dashed) association between waist-to-hip ratio and LDL cholesterol (A) and the reverse (B).** The observed association (red, continuous) was estimated using a similar polynomial approximation for the purpose of comparison, the points showing the mean outcome level at each percentile of the exposure. The hulls surrounding both functions show the 95% confidence range across the distribution.

#### 2.3 BMI

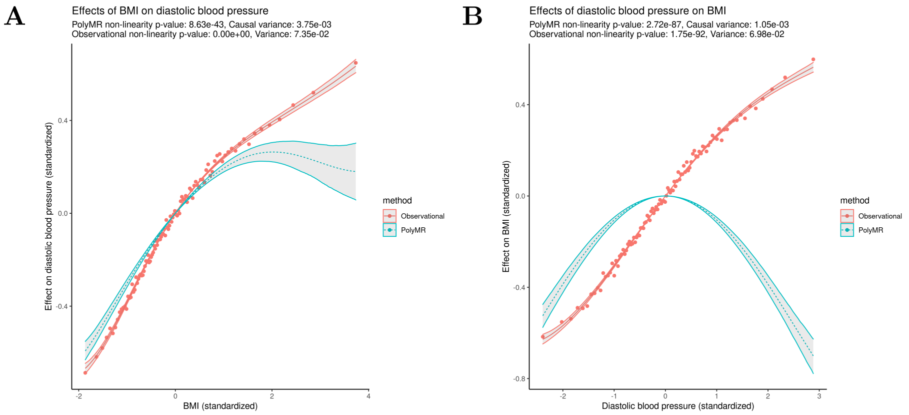

**Supplementary figure 18: The observed (red, continuous) and causal (blue, dashed) association between BMI and diastolic blood pressure (A) and the reverse (B).** The observed association (red, continuous) was estimated using a similar polynomial approximation for the purpose of comparison, the points showing the mean outcome level at each percentile of the exposure. The hulls surrounding both functions show the 95% confidence range across the distribution.

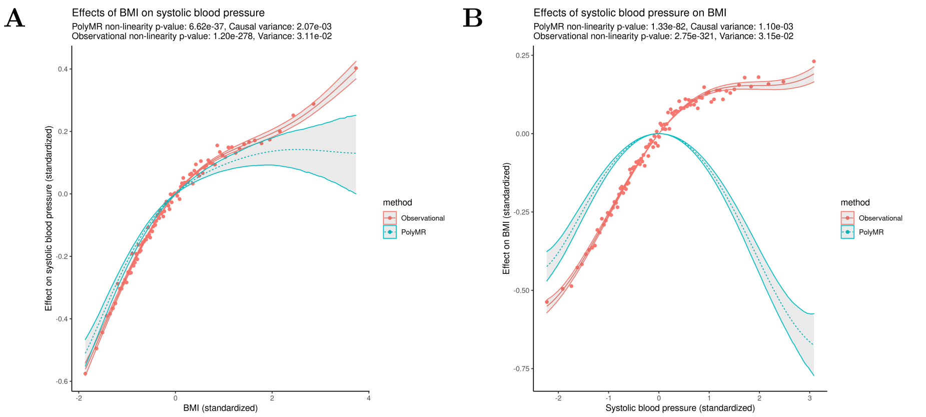

**Supplementary figure 19: The observed (red, continuous) and causal (blue, dashed) association between BMI and systolic blood pressure (A) and the reverse (B).** The observed association (red, continuous) was estimated using a similar polynomial approximation for the purpose of comparison, the points showing the mean outcome level at each percentile of the exposure. The hulls surrounding both functions show the 95% confidence range across the distribution.

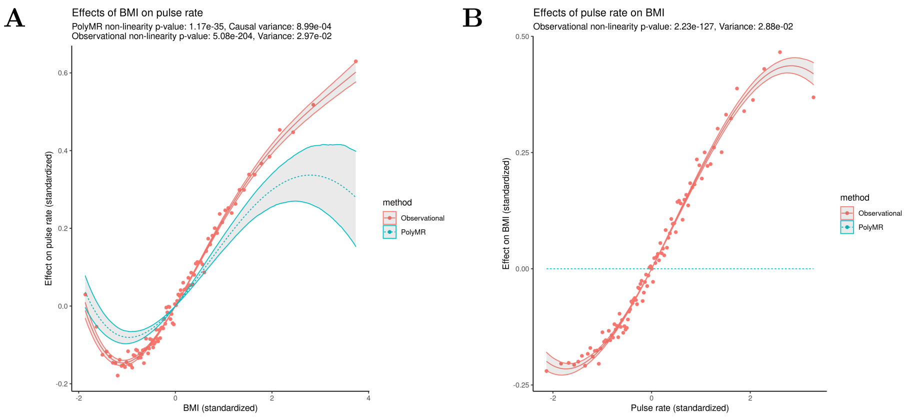

**Supplementary figure 20: The observed (red, continuous) and causal (blue, dashed) association between BMI and pulse rate (A) and the reverse (B).** The observed association (red, continuous) was estimated using a similar polynomial approximation for the purpose of comparison, the points showing the mean outcome level at each percentile of the exposure. The hulls surrounding both functions show the 95% confidence range across the distribution.

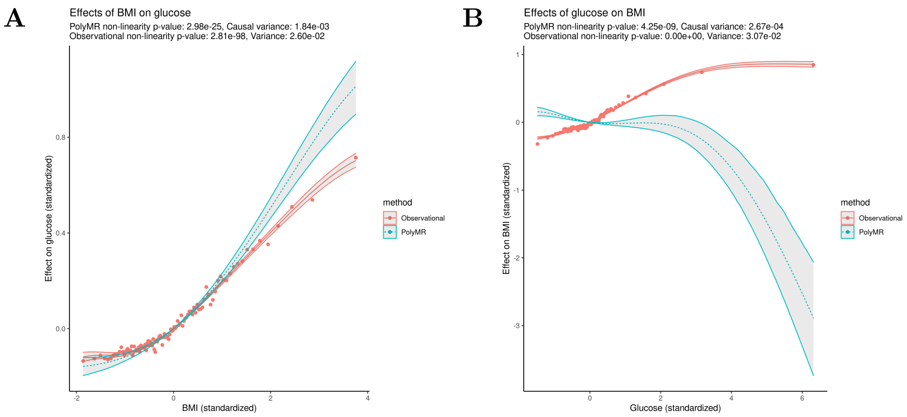

**Supplementary figure 21: The observed (red, continuous) and causal (blue, dashed) association between BMI and glucose (A) and the reverse (B).** The observed association (red, continuous) was estimated using a similar polynomial approximation for the purpose of comparison, the points showing the mean outcome level at each percentile of the exposure. The hulls surrounding both functions show the 95% confidence range across the distribution.

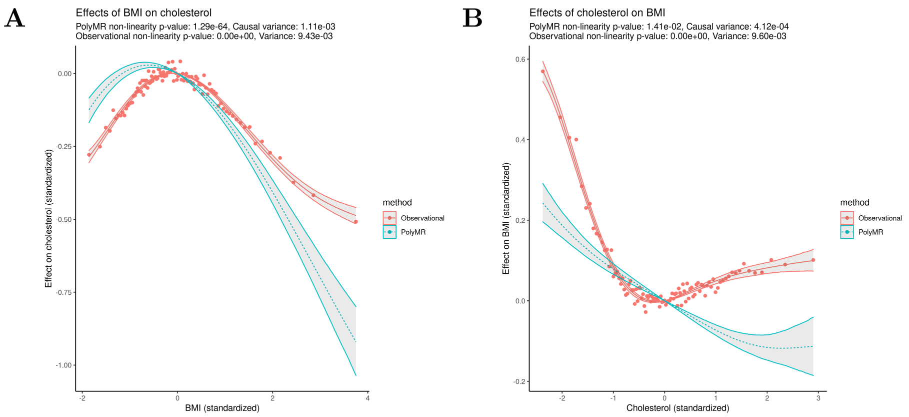

**Supplementary figure 22: The observed (red, continuous) and causal (blue, dashed) association between BMI and total cholesterol (A) and the reverse (B).** The observed association (red, continuous) was estimated using a similar polynomial approximation for the purpose of comparison, the points showing the mean outcome level at each percentile of the exposure. The hulls surrounding both functions show the 95% confidence range across the distribution.

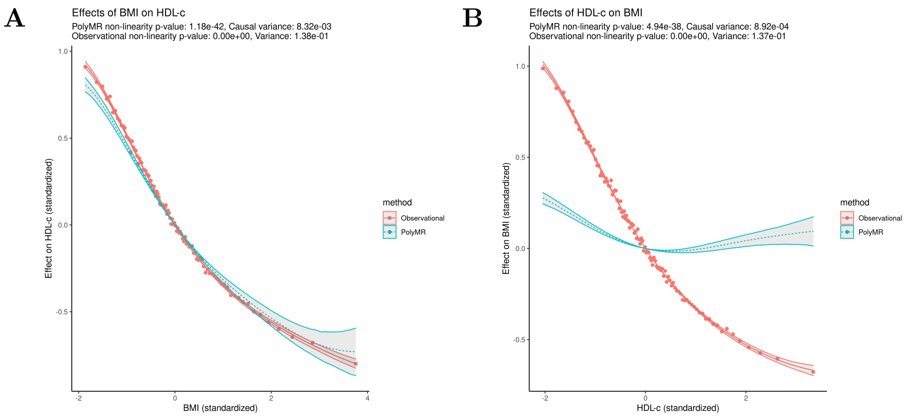

**Supplementary figure 23: The observed (red, continuous) and causal (blue, dashed) association between BMI and HDL cholesterol (A) and the reverse (B).** The observed association (red, continuous) was estimated using a similar polynomial approximation for the purpose of comparison, the points showing the mean outcome level at each percentile of the exposure. The hulls surrounding both functions show the 95% confidence range across the distribution.

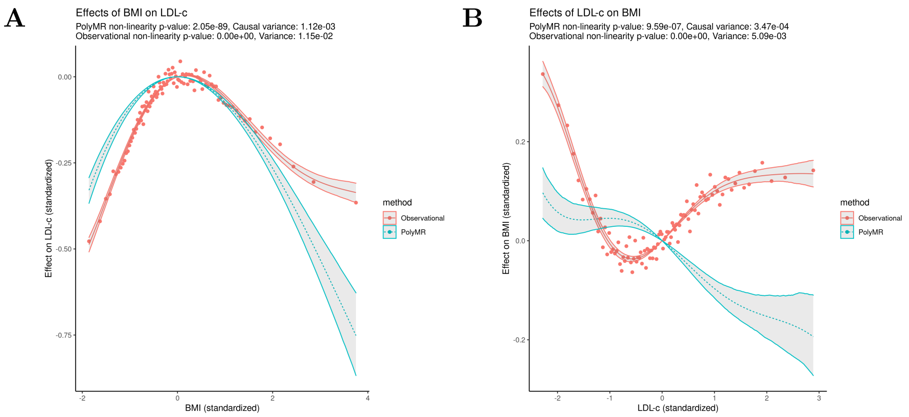

**Supplementary figure 24: The observed (red, continuous) and causal (blue, dashed) association between BMI and LDL cholesterol (A) and the reverse (B).** The observed association (red, continuous) was estimated using a similar polynomial approximation for the purpose of comparison, the points showing the mean outcome level at each percentile of the exposure. The hulls surrounding both functions show the 95% confidence range across the distribution.

#### 2.4 Weight

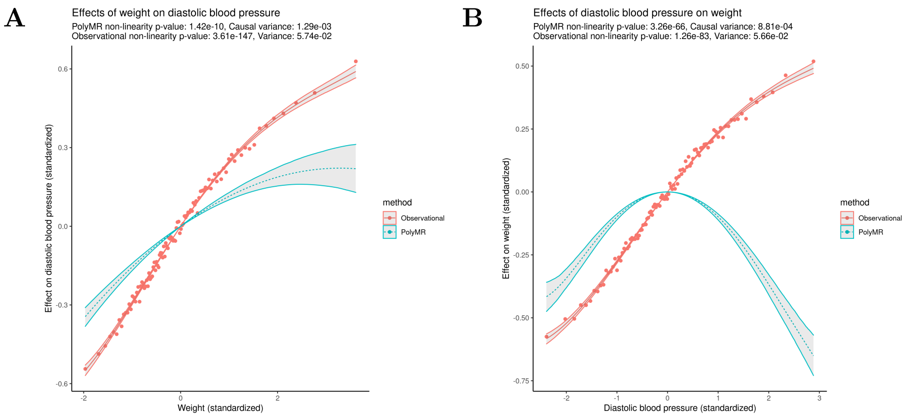

**Supplementary figure 25: The observed (red, continuous) and causal (blue, dashed) association between weight and diastolic blood pressure (A) and the reverse (B).** The observed association (red, continuous) was estimated using a similar polynomial approximation for the purpose of comparison, the points showing the mean outcome level at each percentile of the exposure. The hulls surrounding both functions show the 95% confidence range across the distribution.

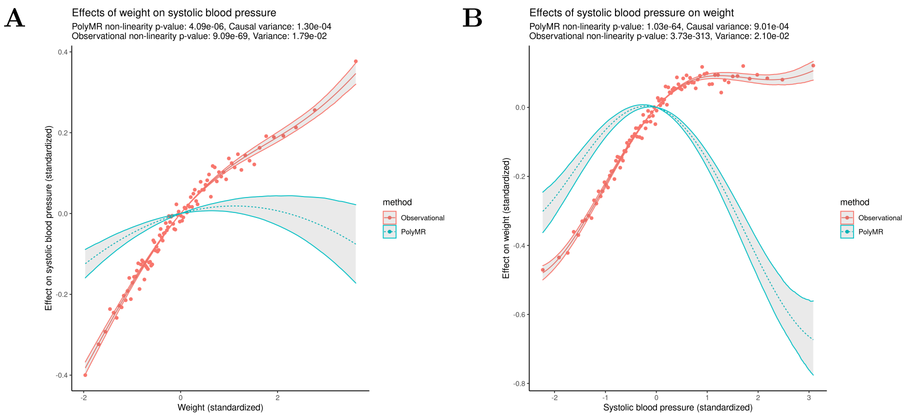

**Supplementary figure 26: The observed (red, continuous) and causal (blue, dashed) association between weight and systolic blood pressure (A) and the reverse (B).** The observed association (red, continuous) was estimated using a similar polynomial approximation for the purpose of comparison, the points showing the mean outcome level at each percentile of the exposure. The hulls surrounding both functions show the 95% confidence range across the distribution.

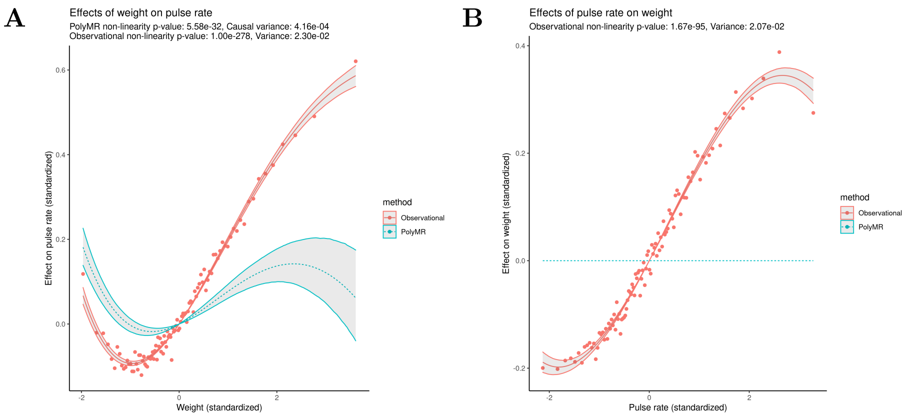

**Supplementary figure 27: The observed (red, continuous) and causal (blue, dashed) association between weight and pulse rate (A) and the reverse (B).** The observed association (red, continuous) was estimated using a similar polynomial approximation for the purpose of comparison, the points showing the mean outcome level at each percentile of the exposure. The hulls surrounding both functions show the 95% confidence range across the distribution.

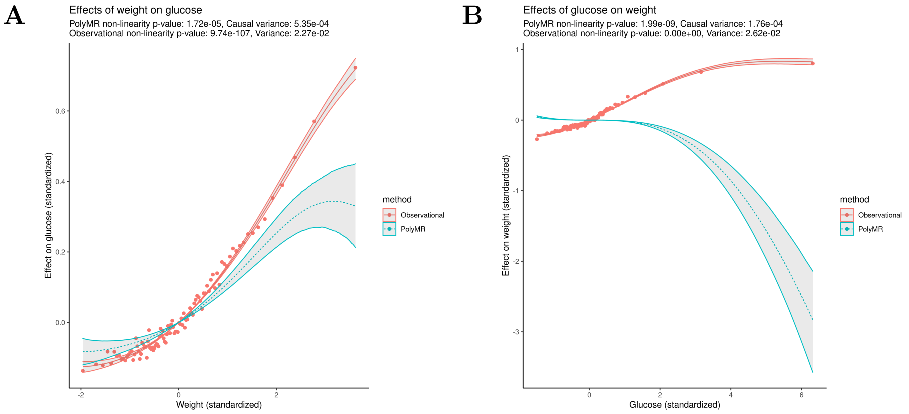

**Supplementary figure 28: The observed (red, continuous) and causal (blue, dashed) association between weight and glucose (A) and the reverse (B).** The observed association (red, continuous) was estimated using a similar polynomial approximation for the purpose of comparison, the points showing the mean outcome level at each percentile of the exposure. The hulls surrounding both functions show the 95% confidence range across the distribution.

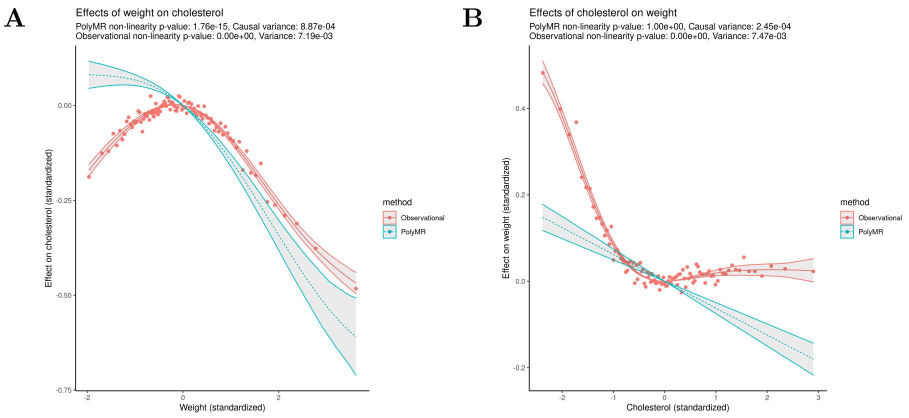

**Supplementary figure 29: The observed (red, continuous) and causal (blue, dashed) association between weight and total cholesterol (A) and the reverse (B).** The observed association (red, continuous) was estimated using a similar polynomial approximation for the purpose of comparison, the points showing the mean outcome level at each percentile of the exposure. The hulls surrounding both functions show the 95% confidence range across the distribution.

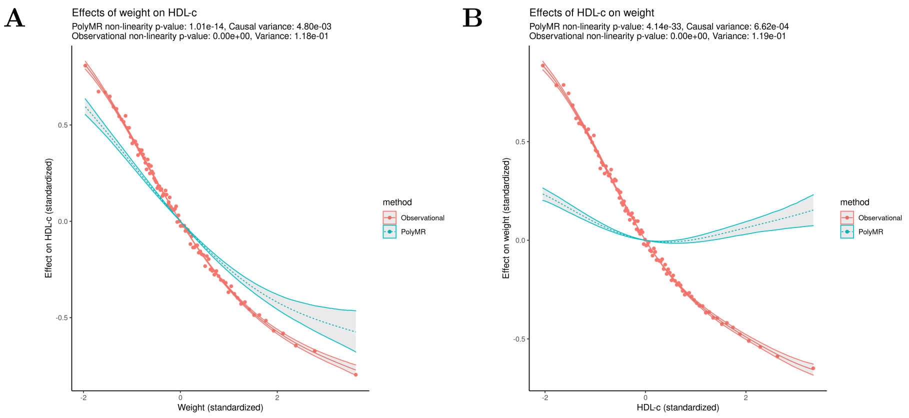

**Supplementary figure 30: The observed (red, continuous) and causal (blue, dashed) association between weight and HDL cholesterol (A) and the reverse (B).** The observed association (red, continuous) was estimated using a similar polynomial approximation for the purpose of comparison, the points showing the mean outcome level at each percentile of the exposure. The hulls surrounding both functions show the 95% confidence range across the distribution.

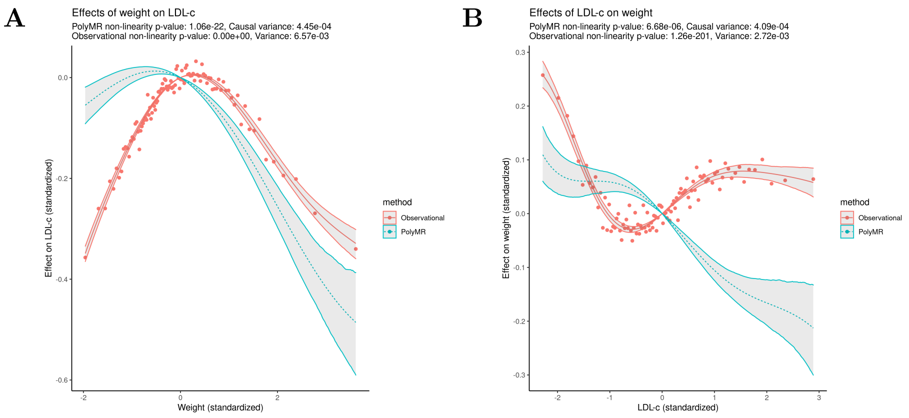

**Supplementary figure 31: The observed (red, continuous) and causal (blue, dashed) association between weight and LDL cholesterol (A) and the reverse (B).** The observed association (red, continuous) was estimated using a similar polynomial approximation for the purpose of comparison, the points showing the mean outcome level at each percentile of the exposure. The hulls surrounding both functions show the 95% confidence range across the distribution.

#### 2.5 Education and life expectancy

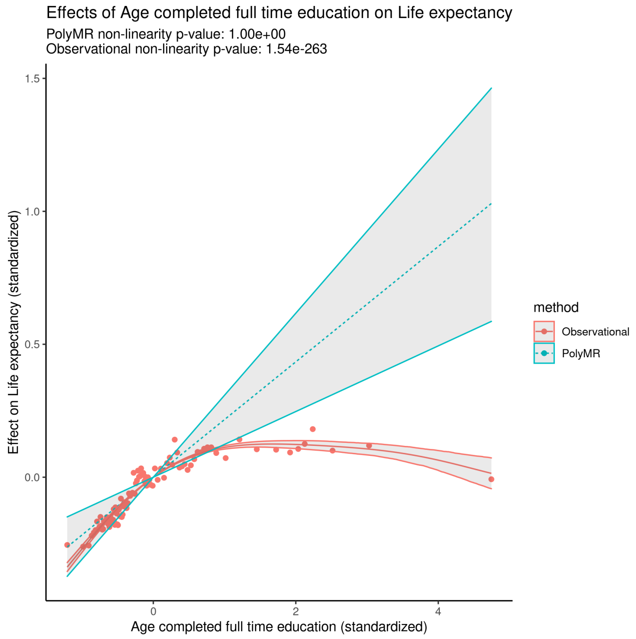

**Supplementary figure 32: The observed (red, continuous) and causal (blue, dashed) association between age completed full-time education and life expectancy.** The observed association (red, continuous) was estimated using a similar polynomial approximation for the purpose of comparison, the points showing the mean outcome level at each percentile of the exposure. The hulls surrounding both functions show the 95% confidence range across the distribution.

### 3 PolyMR-L1

#### 3.1 BMI

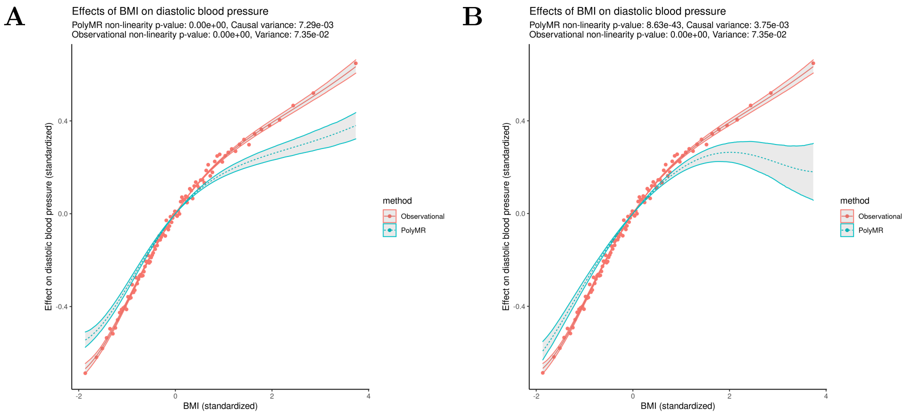

**Supplementary figure 33: The observed (red, continuous) and causal (blue, dashed) association between BMI and diastolic blood pressure, using PolyMR-L1 (accounting for linear confounding alone, A) and PolyMR (B).** The observed association (red, continuous) was estimated using a similar polynomial approximation for the purpose of comparison, the points showing the mean outcome level at each percentile of the exposure. The hulls surrounding both functions show the 95% confidence range across the distribution. Plot B is reproduced here for convenience but is identical to that included above.

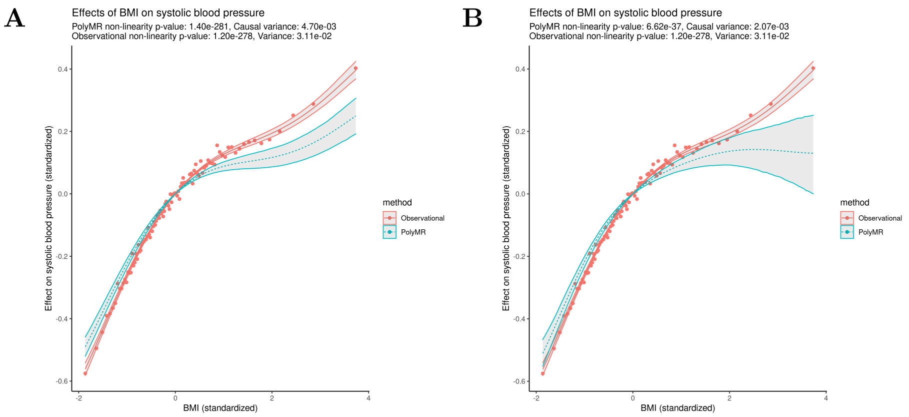

**Supplementary figure 34: The observed (red, continuous) and causal (blue, dashed) association between BMI and systolic blood pressure, using PolyMR-L1 (accounting for linear confounding alone, A) and PolyMR (B).** The observed association (red, continuous) was estimated using a similar polynomial approximation for the purpose of comparison, the points showing the mean outcome level at each percentile of the exposure. The hulls surrounding both functions show the 95% confidence range across the distribution. Plot B is reproduced here for convenience but is identical to that included above.

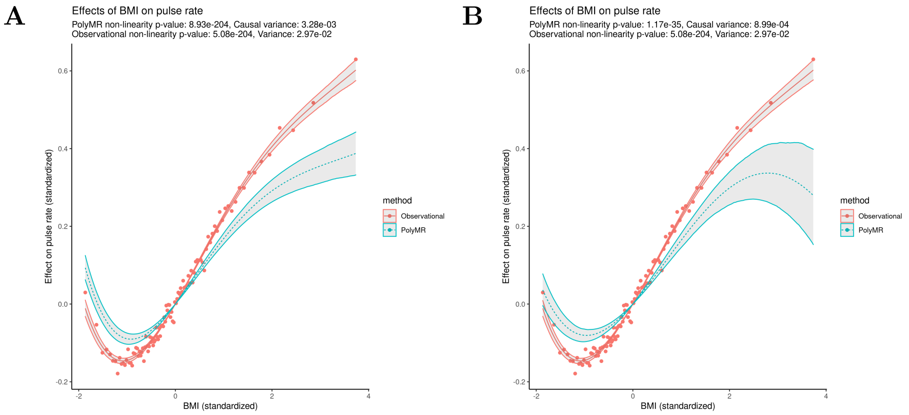

**Supplementary figure 35: The observed (red, continuous) and causal (blue, dashed) association between BMI and pulse rate, using PolyMR-L1 (accounting for linear confounding alone, A) and PolyMR (B).** The observed association (red, continuous) was estimated using a similar polynomial approximation for the purpose of comparison, the points showing the mean outcome level at each percentile of the exposure. The hulls surrounding both functions show the 95% confidence range across the distribution. Plot B is reproduced here for convenience but is identical to that included above.

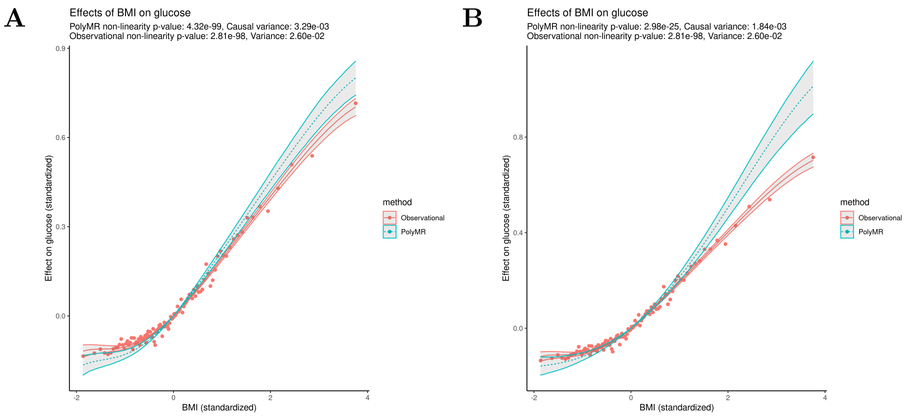

**Supplementary figure 36: The observed (red, continuous) and causal (blue, dashed) association between BMI and glucose, using PolyMR-L1 (accounting for linear confounding alone, A) and PolyMR (B).** The observed association (red, continuous) was estimated using a similar polynomial approximation for the purpose of comparison, the points showing the mean outcome level at each percentile of the exposure. The hulls surrounding both functions show the 95% confidence range across the distribution. Plot B is reproduced here for convenience but is identical to that included above.

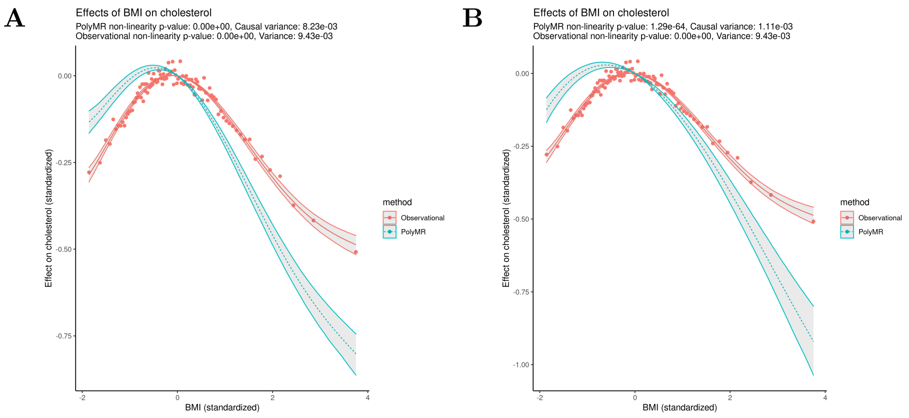

**Supplementary figure 37: The observed (red, continuous) and causal (blue, dashed) association between BMI and total cholesterol, using PolyMR-L1 (accounting for linear confounding alone, A) and PolyMR (B).** The observed association (red, continuous) was estimated using a similar polynomial approximation for the purpose of comparison, the points showing the mean outcome level at each percentile of the exposure. The hulls surrounding both functions show the 95% confidence range across the distribution. Plot B is reproduced here for convenience but is identical to that included above.

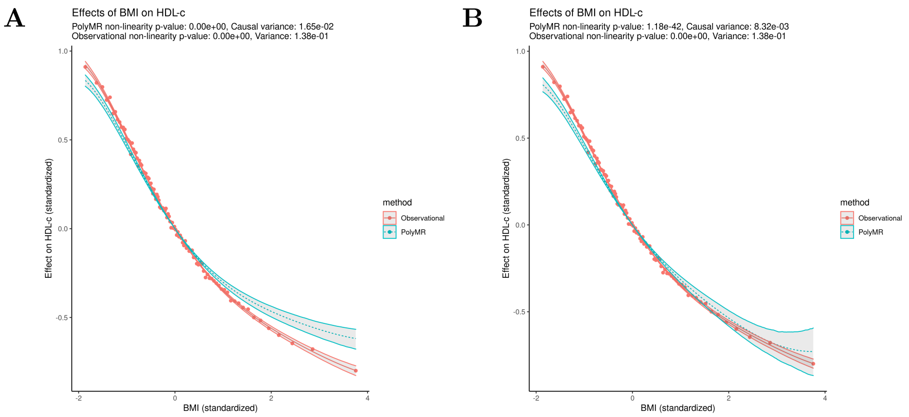

**Supplementary figure 38: The observed (red, continuous) and causal (blue, dashed) association between BMI and HDL cholesterol, using PolyMR-L1 (accounting for linear confounding alone, A) and PolyMR (B).** The observed association (red, continuous) was estimated using a similar polynomial approximation for the purpose of comparison, the points showing the mean outcome level at each percentile of the exposure. The hulls surrounding both functions show the 95% confidence range across the distribution. Plot B is reproduced here for convenience but is identical to that included above.

A

B

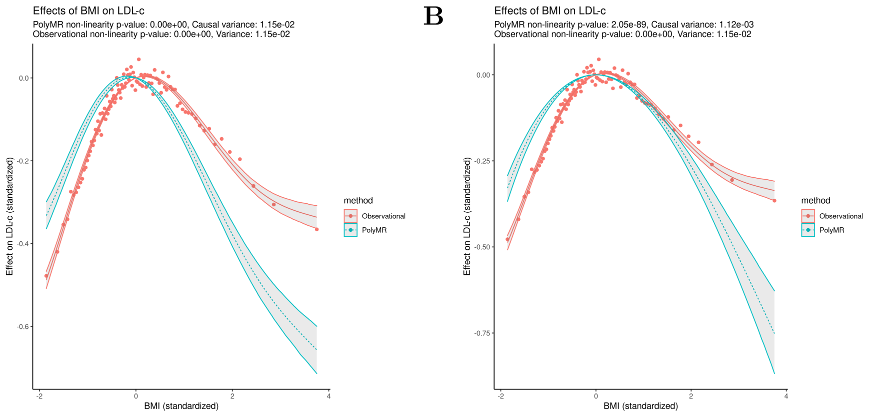

**Supplementary figure 39: The observed (red, continuous) and causal (blue, dashed) association between BMI and LDL cholesterol, using PolyMR-L1 (accounting for linear confounding alone, A) and PolyMR (B).** The observed association (red, continuous) was estimated using a similar polynomial approximation for the purpose of comparison, the points showing the mean outcome level at each percentile of the exposure. The hulls surrounding both functions show the 95% confidence range across the distribution. Plot B is reproduced here for convenience but is identical to that included above.
